# MeSSII interplays the amylose and amylopectin biosynthesis by protein interactions in cassava storage root

**DOI:** 10.1101/2020.03.25.006957

**Authors:** Shutao He, Xiaomeng Hao, Shanshan Wang, Wenzhi Zhou, Qiuxiang Ma, Xinlu Lu, Luonan Chen, Peng Zhang

## Abstract

Starch is a glucose polymer synthesized by green plants for energy storage, and is crucial for plant growth and reproduction. The biosynthesis of starch polysaccharides is mediated by members of the large starch synthase (SS) protein superfamily. Here, we showed that in cassava storage roots, soluble starch synthase II (MeSSII) plays an important role in starch biosynthesis via forming protein complexes with other starch biosynthetic enzymes by directly interacting with MeSSI, MeSBEII and MeISAII. The *MeSSII*-RNAi cassava lines showed increased amylose content and reduced intermediate chain of amylopectin (B1 type) biosynthesis in their storage roots, leading to altered starch physico-chemical properties. Further gel permeation chromatography analysis of starch biosynthetic enzymes between the wild type and *MeSSII*-RNAi lines confirmed the key rule of MeSSII in the organization of heteromeric starch synthetic protein complexes, including the MeSSII-MeSSI-MeGBSSI and MeSBEII-MeSSI-MeSSII-MeISAII-MeISAI complexes. The lack of MeSSII in cassava also reduced the binding capacity of the MeSSI, MeSBEII, MeISAI, and MeISAII to starch granules. Our results show a close coordination between granule-bound starch synthase and amylopectin biosynthetic enzymes, which implies that the processes of amylose synthesis and amylopectin synthesis are directly interrelated. These findings shed light on the key components of the starch biosynthesis machinery in root crops.

**One Sentence Summary:** Here, by focusing on cassava SSII, we elucidated its function and the molecular mechanism by which it chaperones the starch synthase complex to regulate starch biosynthesis in cassava storage roots.

## Introduction

Cassava (*Manihot esculenta* Crantz) accumulates large amounts of starch (up to 32% by fresh weight) in its storage roots even under unfavorable growth conditions in tropical and subtropical regions (Cock 1982). Cassava starch is pure white and contains low levels of fat (0.08–1.54%) and proteins (0.03–0.6%), providing an irreplaceable raw material for bioindustrial applications (Li, Cui et al. 2017). Cassava starch granules are generally round and truncated, ranging in size from 5-40 μm, and the amylose content varies from 15% to 30%, depending on the cultivar and growth conditions (Moorthy 2002). Various modified cassava starch derivatives are widely used in the food, textile, paper manufacturing, pharmaceutical, and bioethanol production industries (Zhu 2015). Despite this, the key enzymes and their regulation of starch biosynthesis in cassava storage roots is largely uncharacterized.

Similar to starches from other plant sources, cassava starch in its native form consists of both amylose and amylopectin. Basically, amylose is a largely linear polymer in which glucosyl residues are joined by α(1→4) bonds, and is an important and interspersed component of starch, but is not crucial for the spatial structure in the formation of starch granules. Amylopectin is composed of a clustered arrangement of crystalline, densely packed glucosyl residues organized as parallel chains, in which linear α-helices are joined by α(1→4) bonds, and interspersed by more disorganized α(1→6) branches at regular intervals of ∼20-30 glucose units. Such organization eventually forms a classic semicrystalline model that determines the stable and insoluble granular structure of starch (Hizukuri 1986). Typically, branched amylopectin chains are classified as A, with a degree of polymerization (DP) of 6-12, B_1_ (DP 13-24), B_2_ (DP 25-36), and B_3+_ (DP ≥37) (Hizukuri 1985). The chain length of cassava starch averages 23.2 DP, and the majority consists of up to 43.7% B_1_ chains; the molecular weight of native cassava starch varies from 1.885 × 10^7^ to 3.323 × 10^7^ g/mol, depending on cultivars and growth conditions (Liu, Zu et al. 2018).

Amylose is synthesized by a single enzyme, granule-bound starch synthase (GBSS), and mutants of cassava lacking GBSS produce amylose-free waxy starches (Ceballos, Sanchez et al. 2007, Zhao, Dufour et al. 2011, Bull, Seung et al. 2018). Recent studies have shown that the activity of GBSS can be regulated by post-translational modifications. It was found that in maize, GBSS is phosphorylated in the starch granule (Grimaud, Rogniaux et al. 2008), which is supported by a recent report of rice endosperm GBSS (Liu, Huang et al. 2013). Liu *et al*. (2013) also noted that GBSS can form oligomers, and ATP and protein kinases increase the degree of oligomerization. In addition, with increasing concentrations of ADP-glucose, GBSS oligomerization increases. However, it is still no solid evidence to support whether GBSS associates with amylopectin synthases directly in the plastids.

The synthesis of amylopectin, which mainly described from grain seeds, requires the coordinated participation of at least 10 different enzymes, including soluble starch synthases (SSs), starch branching enzymes (BEs or SBEs), and debranching enzymes (DBEs). SSs (EC 2.4.1.21) elongate α-1,4-linked linear glucan polymers using ADP-glucose as a substrate. SBEs (EC 2.4.1.18) produce α-1,6-linked branches and are crucial for amylopectin formation because they are the only enzymes that can form branches in amylopectin. DBEs such as isoamylase (ISA; EC 3.2.1.68) and pullulanase (PUL; EC 3.2.1.41) hydrolyze and remove α-1,6-linked branches, which is essential to organize crystalline structure formation in amylopectin (Nakamura 2002, Jeon, Ryoo et al. 2010). In addition, starch phosphorylase (Pho; EC 2.4.1.1) elongates α-1,4-linked linear glucan polymers using glucose 1-phosphate (G1P) as a substrate, and is thought to be involved in the initiation process of starch biosynthesis (Satoh, Shibahara et al. 2008, Jeon, Ryoo et al. 2010). Although GBSSI has an effect on the extra-long amylopectin chains in addition to participating in amylose synthesis (Hanashiro, Itoh et al. 2008), no direct evidence of its interaction with other amylopectin biosynthetic enzymes has been reported. Recently, starches with remarkably altered physico-chemical properties and amylopectin structure have been observed in transgenic cassava in which the expressions of genes for GBSS1 and SBEs have been silenced (Zhao, Dufour et al. 2011, Zhou, Zhao et al. 2020). Nevertheless, the function and regulation of these SSs in cassava remain unclear.

Previous studies have shown that SS can interact with SBE isozymes in wheat, maize, barley, and rice seeds (Tetlow, Wait et al. 2004, Hennen-Bierwagen, Liu et al. 2008, Tetlow, Beisel et al. 2008, Liu, Ahmed et al. 2012, Liu, Romanova et al. 2012, Ahmed, Tetlow et al. 2015, Crofts, Abe et al. 2015). In maize, the trimeric complex composed of SSI, SSIIa, and SBEIIb is one of the best studied, and the formation of the trimer depends on protein phosphorylation (Liu, Ahmed et al. 2012, Liu, Romanova et al. 2012, Makhmoudova, Williams et al. 2014, Mehrpouyan, Menon et al. 2020). Protein-protein interactions have also been found between SBEI and SBEIIb, as well as in protein complexes consisting of SSIII, pyruvate orthophosphate dikinase (PPDK), SBEIIb, and AGPase (Hennen-Bierwagen, Lin et al. 2009, Liu, Makhmoudova et al. 2009). In wheat and maize, Pho1 can interact with SBEI and SBEII (Tetlow, Wait et al. 2004, Subasinghe, Liu et al. 2014). Moreover, functional and mutually synergistic relationships between SBEs and Pho1, and SBEs and SSI, have been observed using recombinant rice starch biosynthetic enzymes *in vitro* (Nakamura, Ono et al. 2012, Abe, Asai et al. 2014, Nakamura, Aihara et al. 2014). In addition, there is also the protein-protein interaction between ISA and a carbohydrate-binding module (CBM)-containing protein, FLO6 (Peng, Wang et al. 2014). Recently, GBSSI has been reported to bind to the surface of starch granules by interacting with the PTST protein (Seung, Soyk et al. 2015), and PTST2 also can interact with SSIV to control starch granule initiation in Arabidopsis leaves (Seung, Boudet et al. 2017).

To date, the majority of studies on multi-enzyme complexes have focused on amylopectin synthesis and are limited to the seeds of major gramineous crop species including wheat, barley, maize, and rice. It remains unknown whether there are similar starch synthase complexes in dicotyledonous species and those with underground storage organs. In this study, we found that in cassava, MeSSII affects starch metabolism by forming protein complexes with other starch biosynthetic enzymes. In particular, in contrast to the current dogma, our results showed a close coordination between granule-bound starch synthase (GBSSI) and amylopectin biosynthetic enzymes, which implies that the processes of amylose synthesis and amylopectin synthesis are directly interrelated.

## Results

### The MeSSII is a member of conserved SSII cluster of starch synthases

Five cassava *SS* genes, which are referred to as *MeSSI, MeSSII, MeSSIII, MeSSIV* and *MeSSV*, were identified after blastp searching the cassava genome using protein sequences of the soluble Arabidopsis starch synthases (SSI to SSV) as queries (Table S1). Phylogenetic analysis of the deduced SS protein sequences from cassava and 15 other plant species showed that the starch synthase protein family can be classified into five categories: the SSI, SSII, SSIII, SSIV, and SSV families (Fig. 1A).

**Figure 1.**
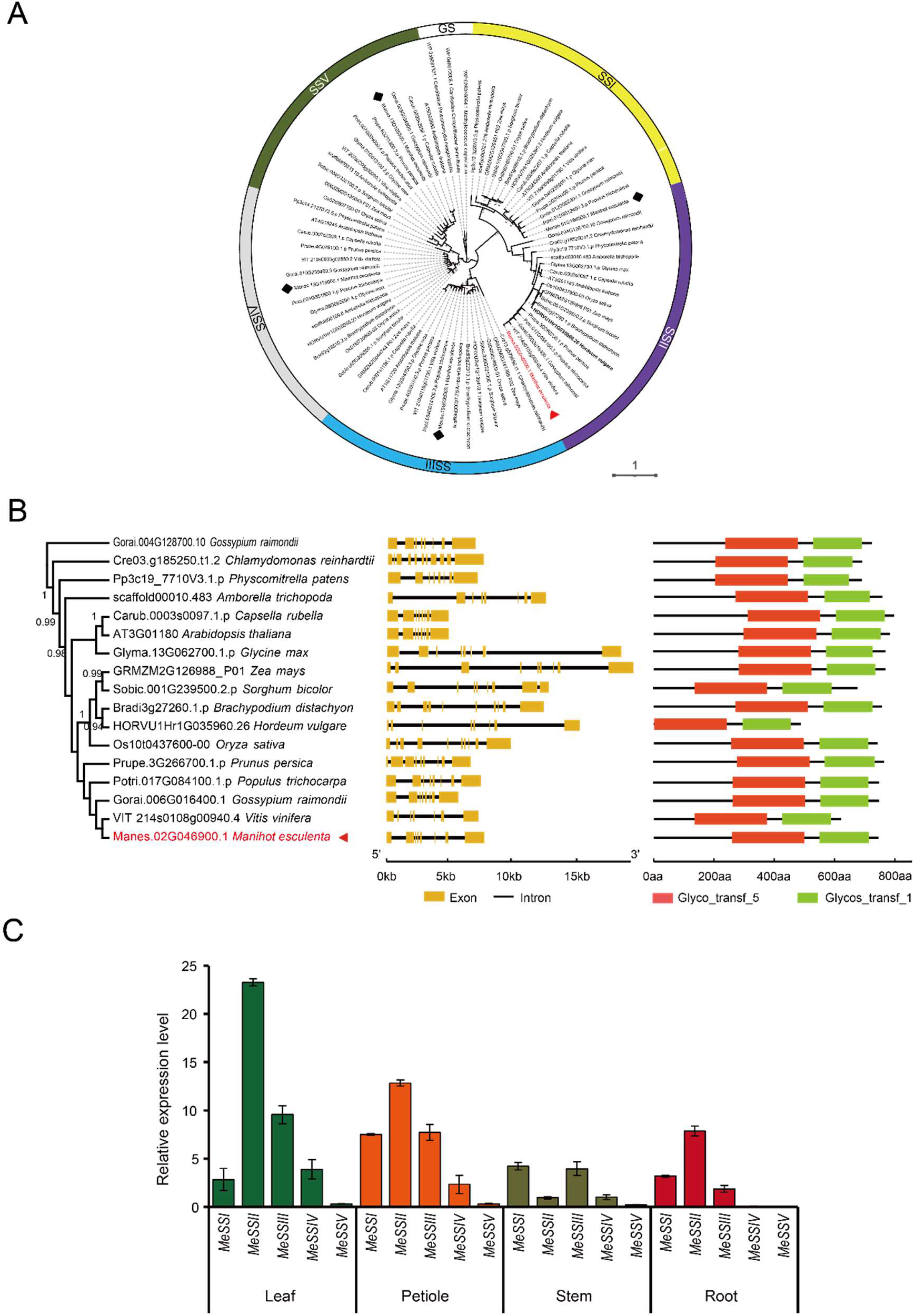
Molecular phylogenies of the plant soluble starch synthase families and transcription of soluble starch synthase genes in cassava. **(A)** Molecular phylogenies of the plant soluble starch synthase families calculated using the maximum likelihood method as implemented in MEGA7. Cassava SSs are marked with a black rhombus (MeSSI, MeSSIII, MessIV and MeSSV) and a red triangle (MeSSII). The closest prokaryotic sequences of At-SS3, At-SS4, and At-SS5 (GS) were included as an outgroup to root the phylogeny. **(B)** Phylogenetic tree, gene structure, and domain analyses of soluble starch synthase II (SSII) proteins. The phylogenetic tree of SSIIs constructed with MEGA using the maximum likelihood method is shown in the left panel. Exon-intron structures of *SSII* genes are shown in the middle panel. Conserved domains annotated using the Pfam database are shown on the right panel. Glyco_transf_1, Glycosyl transferases group 1; Glyco_transf_5, starch synthase catalytic domain. The numbers on the nodes show the bootstrap values. **(C)** mRNA levels of transcribed soluble starch synthase genes in different cassava organs quantified by qRT-PCR assays.

To obtain further insights into the possible structural evolution of the *SSII* genes, the exon-intron structures and conserved domains in these *SSII* genes were visualized using the Gene Structure Display Server (GSDS; http://gsds.cbi.pku.edu.cn/) as shown in Fig. 1B. The numbers of exons varied between 7 and 11. The exon positions also varied between the genes. Although there were various exon-intron patterns present in these species, the conserved domains were found to be highly similar (Fig. 1B). In accordance with previous observations (Liu, Yu et al. 2015, Helle, Bray et al. 2018), all of the deduced SSII proteins were predicted to have Glycos_transf_1 domain and Glyco_transf_5 domain, which are the typical pattern of conserved domains found in the SSII family proteins.

### Down-regulated *MeSSII* expression leads to reduced plant growth and biomass in cassava

To further investigate the function of MeSSII, we first characterized the tissue-specific gene expression patterns of cassava starch synthase genes in the five SS families (Fig. 1C). Among the 5 cassava starch synthase genes, *MeSSV* was almost undetectable in all tissues. In storage roots, *MeSSII* had the highest transcript level among these soluble starch synthase genes, which indicates that *MeSSII* might play an important role in starch metabolism.

A 200 bp conserved DNA sequence of cassava *MeSSII* gene (base pairs 978-1177), which encodes the starch synthase catalytic domain, was used as the target sequence for RNAi. After primary growth and phenotypic screening, three *MeSSII*-RNAi transgenic (SSC) plant lines that showed normal growth and development, together with one empty vector (EV) line, were chosen for further study. Single transgene integrations were detected in all transgenic lines by Southern blot analysis using an *HPT* DIG-labeled probe following digestion of genomic DNA with either *Hin*dIII (Fig. 2A, upper panel) or *Eco*RI (Fig. 2A, lower panel). The results of qRT-PCR (Fig. 2B) and Western blotting analyses using antibody against MeSSII (Fig. 2C) suggested that the expression of *MeSSII* in the storage roots of the SSC lines was significantly decreased compared to the wild type (WT) and EV controls.

**Figure 2.**
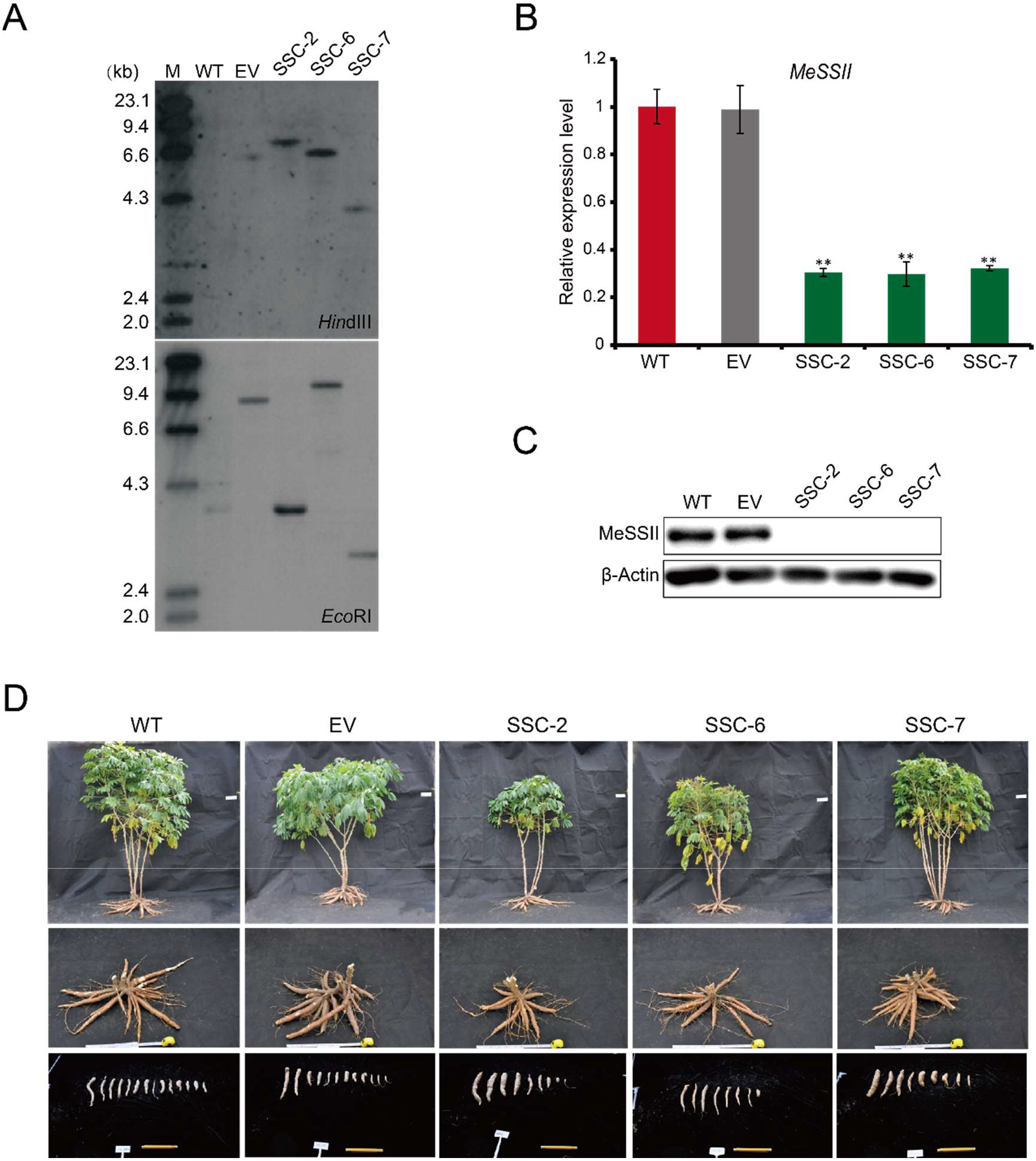
Repression of MeSSII expression affects phenotypes of field-grown cassava. **(A)** Integration patterns of transgenes in SSC transgenic cassava plant lines determined by Southern blotting analysis. M, DIG-labeled molecular marker; WT, wild type control; EV, empty vector plasmid 35::RNAi; SSC-x, plant lines transformed with 35S::MeSSII-RNAi. **(B-C)** mRNA (B) and protein (C) levels of *MeSSII* in storage roots of wild type (WT), empty vector (EV) and *MeSSII*-RNAi (SSC) transgenic plants determined by qRT-PCR and Western blotting using a specific antibody against MeSSII, respectively. The actin protein was used as a loading control. **(D)** Canopy architecture (upper panel), attached storage roots (middle panel), and harvested storage roots (lower panel) of the SSC transgenic plants in comparison with the wild type (WT) and empty vector (EV) plants.

After six months of growth in the field, phenotypic measurements that included plant height, fresh-weight biomass, root number, root length, and root diameter were performed on the WT and transgenic lines. Plant heights ranged from 160.8 cm to 166.6 cm for the SSC lines, which were shorter than both WT and EV line plants (Fig. 2D and Supplementary Fig. S1A). The biomass per plant ranged from 1.39 kg to 2.15 kg for the SSC lines (Fig. 2D and Supplementary Fig. S1B), which was considerably less than the average fresh weight of the WT (4.33 kg) and EV (4.01 kg) plants. The average storage root length was 23.42 cm for WT and 23.33 cm for EV, and it ranged from 19.41 cm to 21.26 cm for the SSC lines (Fig. 2D and Supplementary Fig. S1C). Average root diameter was 2.67 cm for WT, 2.64 cm for EV, and 2.52 cm to 2.88 cm for the SSC lines (Fig. 2D and Supplementary Fig. S1D). The average root number per plant varied from 11.67 to 14 for the SSC lines (Fig. 2D and Supplementary Fig. S1D), which was dramatically less than the numbers for WT (25.1) and EV (25) plants. Thus, for the SSC transgenic lines, total biomass was reduced due to retarded growth of both the aerial and subterranean organs.

### MeSSII influences starch properties in cassava storage roots

Microscopic analysis of storage root starch granule morphology by SEM and TEM did not show obvious differences between the WT and SSC transgenic lines. The starch in both the WT and SSC lines consisted of a mixture of round, truncated, and dome-shaped granules (Fig. 3A and Supplementary Fig. S2). The starch granule size distributions in the SSC transgenic lines differed from those of the WT and EV. The SSC lines exhibited a broader granule size distribution compared to the WT and EV, and contained a greater number of larger starch granules which were largely responsible for the increased average granule size in the SSC lines (Fig. 3B).

**Figure 3.**
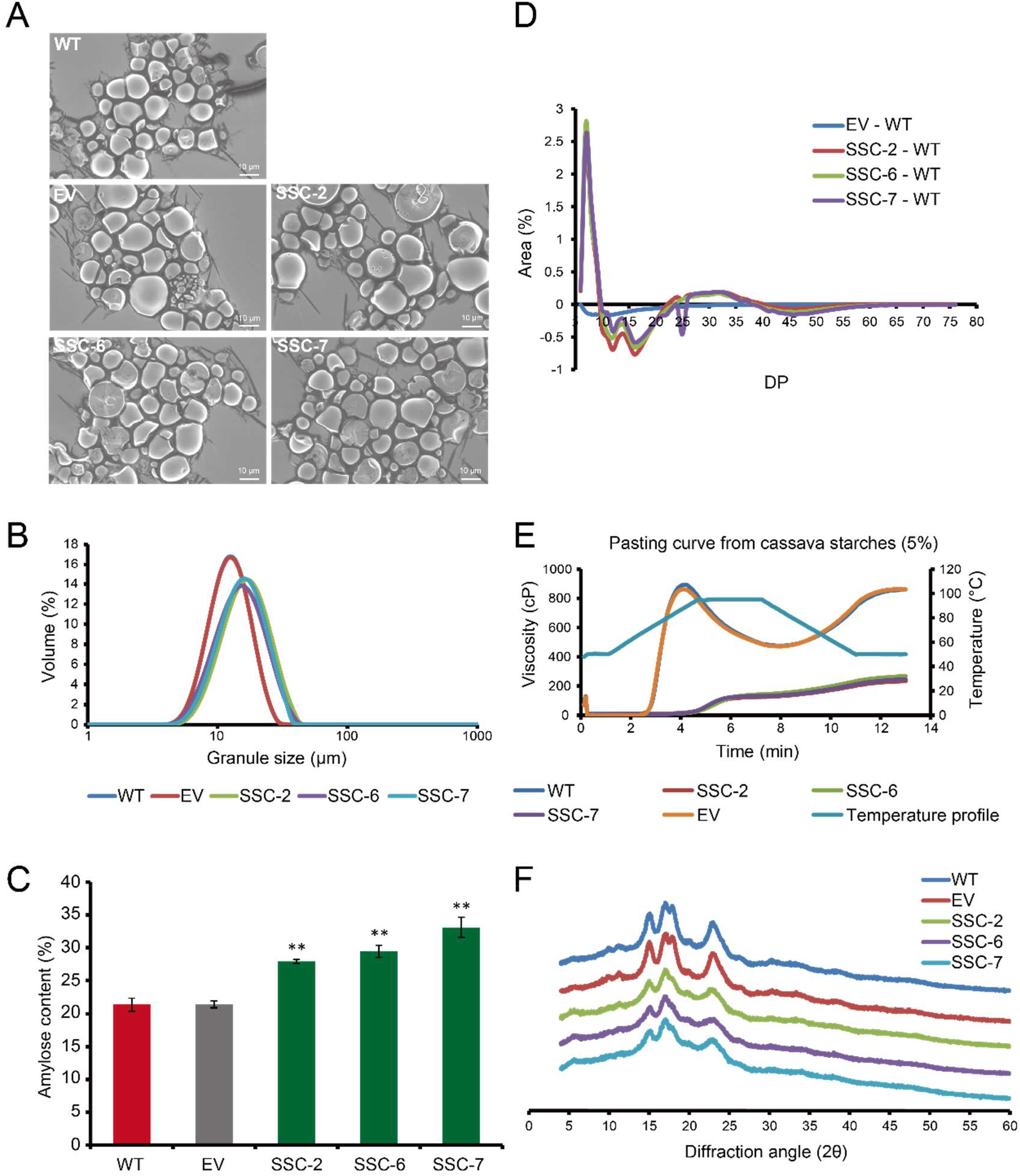
Physico-chemical properties of storage root starches in cassava. **(A, B)** Scanning electron microscopy (A) and starch granule size distribution (B) of starch granules from wild type (WT), empty vector (EV), and SSC transgenic cassava plants. **(C-E)** Amylose content (C), Chain length distribution (D), Rapid Visco Analyser (RVA) pasting profiles (E), and X-ray diffractograms (F) of storage starches isolated from plants of WT, EV, and SSC transgenic plant lines. * and ** indicate a significant difference compared to WT at *P* < 0.05 and < 0.01, respectively, determined by Student’s *t*-test.

The relative proportion of amylose was dramatically increased in the SSC lines compared with starch from the WT and EV (Fig. 3C). In the SSC lines, the number of chains with DP 6-9 and DP 24-40 increased, while the number of chains with DP 10-23 decreased (Fig. 3D). The chain length distribution (CLD) differential patterns in the SSC lines indicated that MeSSII could play a key role in the synthesis of short to intermediate length amylopectin chains (DP 10-23) in cassava storage roots.

RVA analysis showed that root starches from the SSC lines exhibited different pasting patterns compared to WT and EV starches (Fig. 3E). The pasting temperature (PT) of WT starch was 71.05°C, while all three SSC starches showed increased PTs (Table S2). Overall, the values for peak viscosity (PV), hot paste viscosity (HV), breakdown (BD), final viscosity (FV), and setback (SB) were significantly lower in the SSC lines than in starches from the WT and EV lines. The DSC (differential scanning calorimeter) parameters of the starch samples were determined in order to evaluate the gelatinization properties (Table S3). Starch isolated from WT roots gelatinized with a temperature range of 56.26°C (*T*_*o*_) to 78.21°C (*T*_*c*_) and an enthalpy (Δ*H*) of 11 J/g. The gelatinization temperatures (*T*_*o*_, *T*_*p*,_ and *T*_*c*_) and enthalpies (Δ*H*) were decreased in starch isolated from the SSC transgenic lines, and showed clear differences when compared with starches from the WT and EV lines.

X-ray diffraction (XRD) analysis of normal cassava storage starch showed typical A-type crystals with one doublet around 17° (2*θ*) and two singlets predominantly around 15° and 23° (2*θ*) (Fig. 3F) (Zhao, Dufour et al. 2011). Compared to storage root starch from WT and EV plants, SSC storage starches underwent pronounced changes in their diffraction patterns, including replacement of a doublet with a singlet emerging around 17° (2*θ*). The peak intensities around 15°, 17°, and 23° (2*θ*) of SSC starches were weaker than that of WT and EV starches.

### No obvious alternations in protein levels of six key starch biosynthetic enzymes

First, we obtained polyclonal antibodies specific for cassava MeSSI, MeSSII, MeSBEI, MeSBEII, MeISAI, MeISAII and MeGBSSI by means of recombinant protein immunization. Genetic tests to verify the specificity of each antiserum by using RNAi transgenic plants including SSC-6, *sbe1* (Zhou, Zhao et al. 2020), *sbe2* (Zhou, Zhao et al. 2020) and *gbss1* (Zhao, Dufour et al. 2011), which inhibit expression levels of *MeSSII, MeSBEI, MeSBEII* and *MeGBSSI*, respectively. The results demonstrate conclusively that all these antibodies had good specificity and could specifically recognize the corresponding proteins (Supplementary Fig. S3).

Subsequently, the above antibodies were used to analyze the expression and solubility of starch biosynthetic enzymes from cassava mature roots (diameter > 2.5 cm) by immunoblotting. Proteins were extracted using a denaturing buffer which also gelatinizes the starch to enable extraction of granule-bound proteins. The results showed that all of these enzymes (MeSSI, MeSSII, MeSBEI, MeSBEII, MeISAI, MeISAII, and MeGBSSI) can be detected in the soluble part (Supplementary Fig. S4). Generally, MeGBSSI binds to starch granules tightly and is not soluble, but in our analysis, 51% of the MeGBSSI could be detected in the soluble fraction in the cassava storage roots.

To further uncover how MeSSII reduction affects starch properties, the relative mRNA levels of other six key starch biosynthetic genes were determined by qRT-PCR (Supplementary Fig. S5A). Although only *MeSSI* and *MeSBEII* transcription was found to be up-regulated in the SSC lines, all six enzymes showed no significant differences at protein levels in the SSC lines compared to the WT and EV (Supplementary Fig. S5B). These results indicate that the phenotypic differences between the transgenic SSC lines and WT are not due to the expression of these six starch biosynthetic genes.

### MeSSII forms protein complexes with other starch biosynthetic enzymes

To illustrate the molecular weight distribution patterns of seven key starch biosynthetic enzymes (MeSSI, MeSSII, MeSBEI, MeSBEII, MeISAI, MeISAII and MeGBSSI), soluble proteins extracted from mature cassava storage roots were fractionated using gel permeation chromatography (GPC) and blue native polyacrylamide gel electrophoresis (BN-PAGE). All seven enzymes were detected in the higher molecular weight fraction (Fig. 4A, B), which is consistent with the notion that they can form protein complexes.

**Figure 4.**
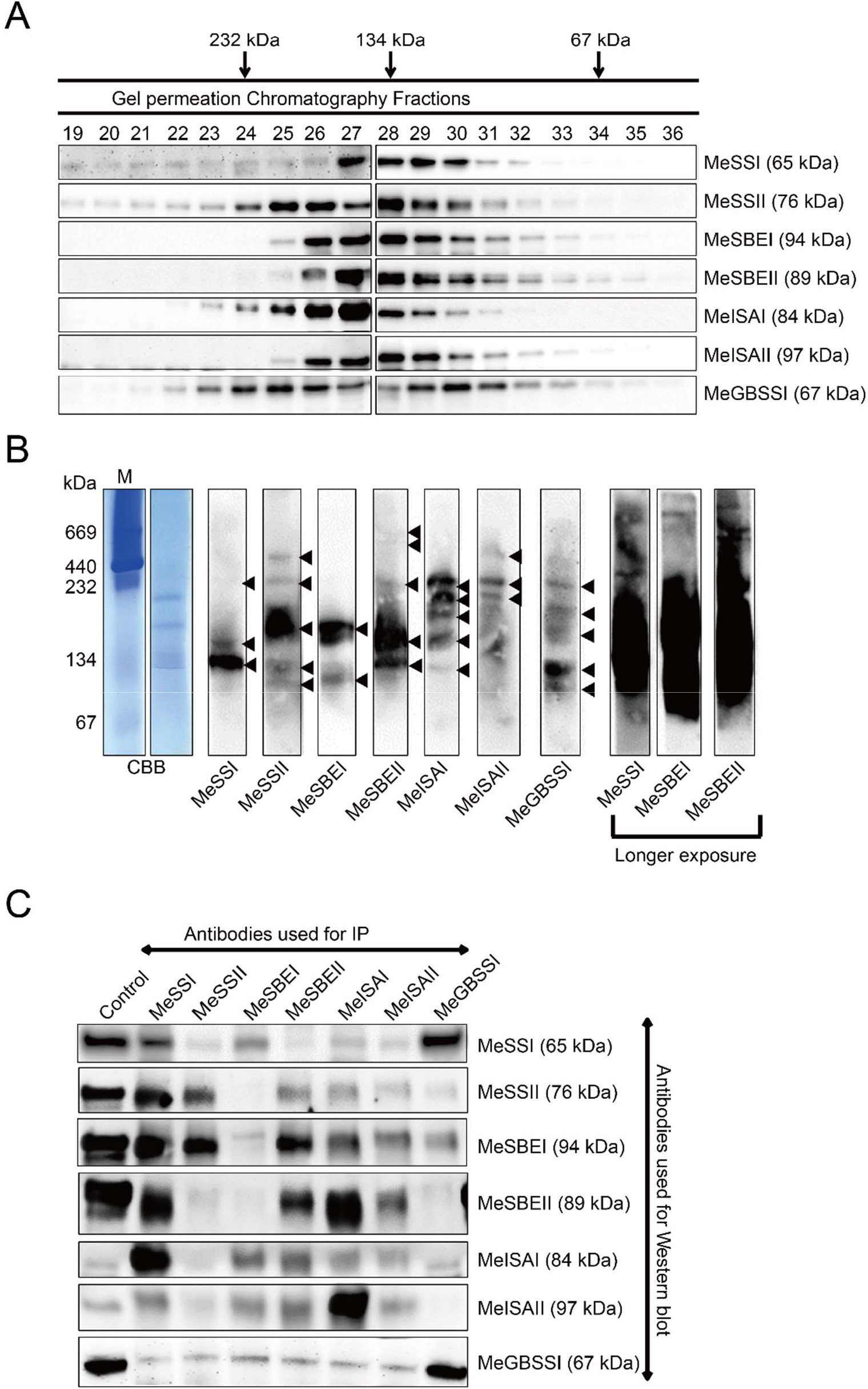
Protein-protein interactions among starch biosynthetic enzymes of cassava. **(A)** GPC analyses of proteins extracted from mature storage roots of wild type (WT) cassava. Proteins present in the indicated fractions from the GPC column were separated by SDS-PAGE and then probed with each specific antibody in immunoblot analyses. The peak elution volumes of molecular mass standards analyzed using the same GPC protocol are indicated at the top. Monomeric molecular weights of each enzyme are indicated on the right. **(B)** Blue native (BN)-PAGE analyses of starch biosynthetic enzyme complexes from mature storage roots of wild type (WT) cassava. Soluble protein extracts were separated by native-PAGE and then stained with Coomassie brilliant blue (CBB) or probed with specific antibodies in immunoblot analyses. Arrowheads indicate the presence of polypeptides recognized by the antibodies. **(C)** Co-immunoprecipitation (Co-IP) analyses of cassava starch biosynthetic enzymes. Soluble proteins from mature storage roots were immunoprecipitated using the corresponding specific antibodies. Soluble proteins from WT cassava storage roots were used as a positive control. The antibodies used for Western blotting are indicated on the right.

To investigate possible interacting partners among these starch biosynthetic isozymes, immunoprecipitation was performed by using soluble cassava storage root protein extracts and isozyme-specific antibodies (Fig. 4C). The results demonstrated that the pairwise protein interactions obtained by reciprocal co-immunoprecipitation includes MeSSI-MeSSII, MeSSI-MeSBEI, MeSSI-MeSBEII, MeSSI-MeISAI, MeSSI-MeISAII, MeGBSSI-MeSSI, MeSSII-MeSBEII, MeSSII-MeGBSSI, MeSBEI-MeISAI, MeSBEI-MeISAII, MeSBEI-MeGBAII, MeSBEII-MeISAII, MeISAI-MeISAII, MeISAI-MeISAII, MeISAI-MeISAII, and MeISAI-MeISAII. Western blot signals obtained from only one side of the co-immunoprecipitation included AbMeSSII-MeSBEI, AbMeSSII-MeISAI, AbMeSSII-MeISAII, AbMeSBEI-MeSBEII, and AbMeISAII-MeGBSSI (first acronym, antibody used for immunoprecipitation; second acronym, isozyme detected by Western blotting).

To further explore the components of the starch biosynthetic isozyme complexes in mature cassava storage roots, protein complexes immunoprecipitated by each antibody were analyzed separately by LC-MS/MS, and the identified proteins are summarized in Table S4. The results showed that MeSSI, MeSSII, MeSBEI, MeSBEII, MeISAI, MeISAII, and MeGBSSI can be co-immunoprecipitated by each antibody, indicating that these enzymes form protein complexes through specific protein-protein interactions, further confirming the Co-IP experiment results (Fig. 4C). In addition, many other proteins were also detected. Among these proteins, the 14-3-3 protein was co-immunoprecipitated by each antibody. Starch phosphorylase (MePho) can be co-immunoprecipitated with MeSBEI and MeSBEII antibodies, which is consistent with previous reports showing that Pho can interact with SBEI and SBEII (Tetlow, Wait et al. 2004, Subasinghe, Liu et al. 2014). MePho is also co-immunoprecipitated by antibodies against MeSSI, MeSSII, MeISAI, MeISAII, and MeGBSSI. Pyruvate orthophosphate dikinase (MePPDK) can be co-immunoprecipitated with MeSSII and MeGBSSI, which is consistent with previous reports that PPDK can form protein complexes with starch biosynthetic enzymes (Hennen-Bierwagen, Lin et al. 2009). Furthermore, each antibody can co-immunoprecipitate with MeAGPase and MeSuS.

To investigate direct protein-protein interaction patterns among these seven enzymes, we performed yeast-two-hybrid (Y2H) assays. We found that MeSSI, MeSSII, and MeSBEII can interact with one another (Fig. 5A). MeSSII can interact with itself and MeISAII directly, and the interaction between MeSSII and MeISAII was a very interesting finding, indicating that SSs, SBEs and ISAs may assemble into multi-enzymes complexes to perform their functions. Furthermore, we found that MeGBSSI could interact with itself and was also capable of interacting with MeSSI and MeISAI (Fig. 5A). In addition, we detected strong interactions between MeISAI and MeISAII, and MeISAII with itself.

**Figure 5.**
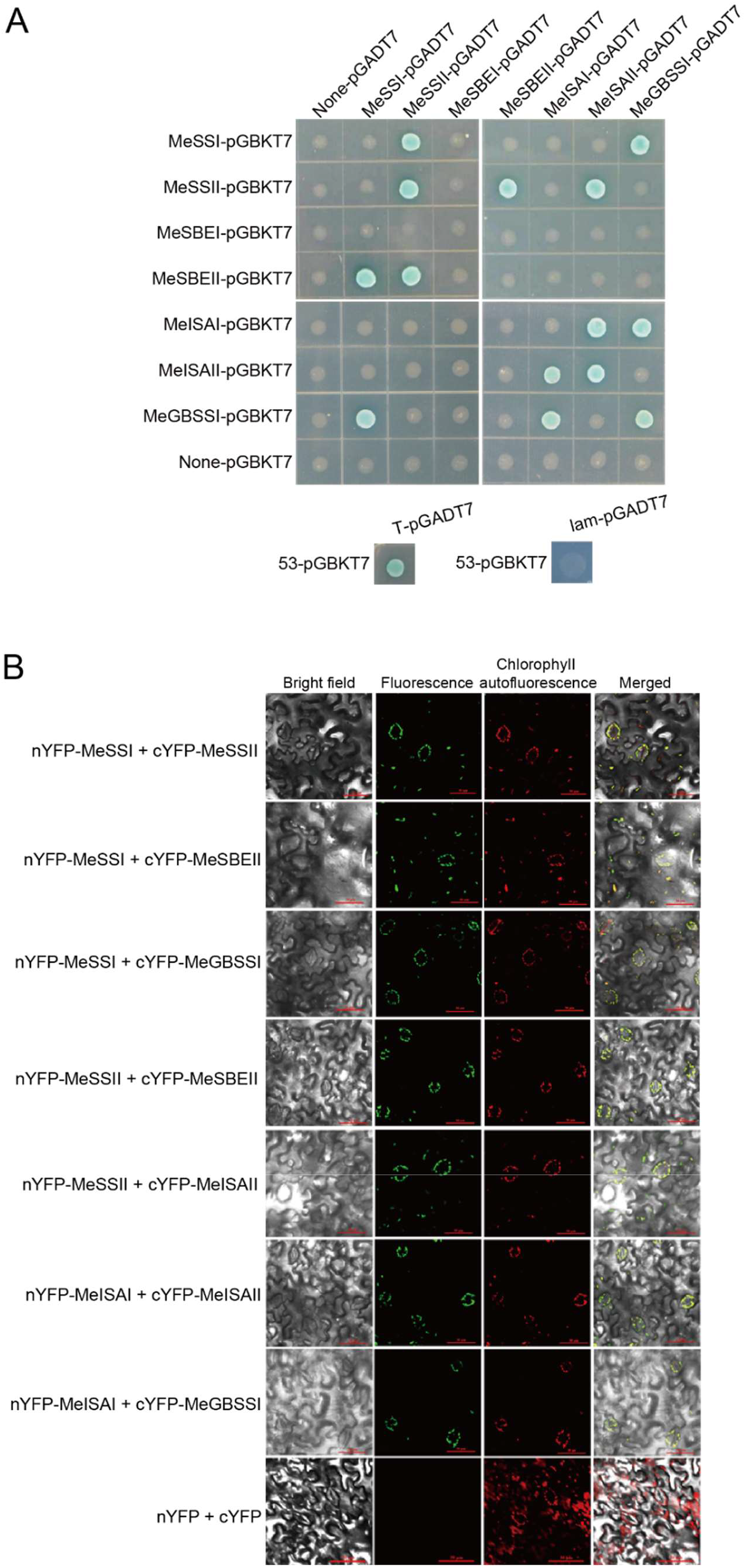
*In vivo* protein-protein interactions between starch biosynthetic enzymes of cassava. **(B)** Yeast two-hybrid analyses. Empty vectors (None-pGBKT7 and None-pGADT7) were used as the negative control. 53-pGBKT7/T-pGADT7 was used as the positive control, and 53-pGBKT7/lam-pGADT7 was used as the negative control. **(B)** Bimolecular fluorescence complementation (BiFC) assays. The empty vectors (nYFP+cYFP) were used as a negative control.

The Y2H results were validated by testing these interactions in bimolecular fluorescence complementation (BiFC) experiments. The cDNAs encoding the above seven enzymes were cloned into BiFC vectors to generate protein fusions with either the N-terminal half of YFP (nYFP) or the C-terminal half of YFP (cYFP). Vector combinations of nYFP-MeSSI+cYFP-MeSSII, nYFP-MeSSI+cYFP-MeSBEII, nYFP-MeSSI+cYFP-MeGBSSI, nYFP-MeSSII+cYFP-MeSBEII, nYFP-MeSSII+cYFP-MeISAII, nYFP-MeISAI+cYFP-MeISAII, and nYFP-MeISAI+cYFP-MeGBSSI were transiently co-expressed in *N. benthamiana* leaves, and strong YFP fluorescence signals were detected in the chloroplasts (Fig. 5B and Supplementary Fig. S6), which were consistent with the Y2H results (Fig. 5A).

### MeSSII influences GPC mobility and granule-bound ability of starch biosynthetic enzymes

To investigate whether the absence of MeSSII would alter the GPC mobility of other members of the complex, soluble proteins extracted from mature roots of the WT and SSC lines were fractionated using GPC (Fig. 6A). MeSSI and MeSBEII mobility in the GPC column was significantly affected by the absence of MeSSII. In the WT protein extract, MeSBEII was not detected in the 232 kDa fraction, and the amount of MeSSI present in the 232 kDa fraction was reduced compared with the SSC lines. No obvious changes in the GPC mobility of other proteins (MeSBEI, MeISAI, MeISAII, and MeGBSSI) were detected in the SSC lines compared with WT. The fact that MeSSI and MeSBEII exhibited altered GPC elution volumes when MeSSII was depleted, compared with WT, is consistent with the proposal that the three proteins associate with one another in the same quaternary structure.

**Figure 6.**
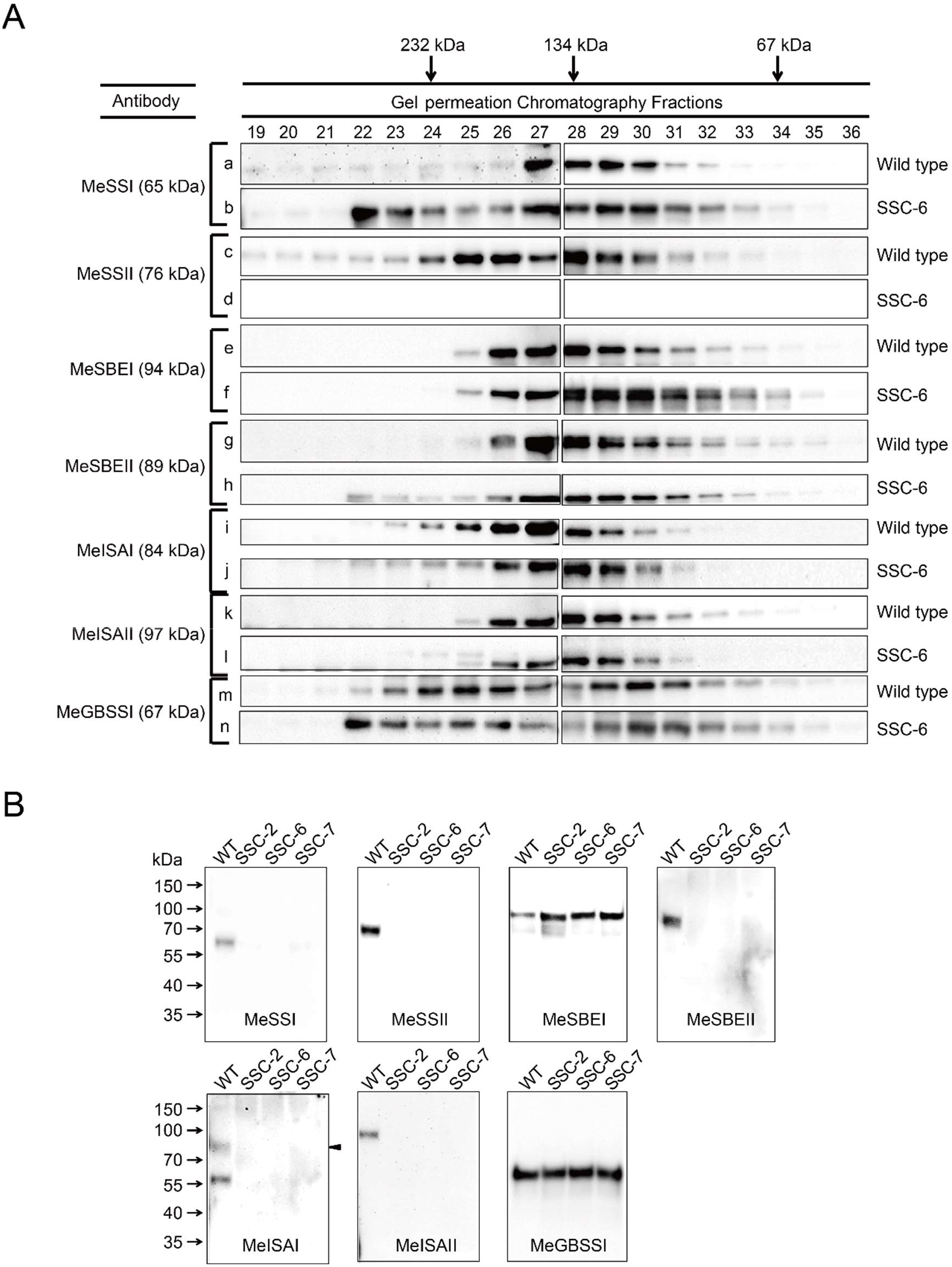
Characterization of protein interaction patterns and starch granule-bound proteins in storage roots between WT and SSC transgenic cassava plants. **(A)** GPC analyses of proteins extracted from mature storage roots of wild type (WT) and SSC transgenic cassava plants. Proteins present in the indicated fractions from the GPC column were separated by SDS-PAGE and then probed with specific antibodies in immunoblot analyses. The peak elution volumes of molecular mass standards analyzed using the same GPC protocol are indicated at the top. The monomeric molecular weights of each enzyme are indicated on the left. Material names are indicated on the right. **(B)** Analysis of starch granule-bound proteins in storage roots of WT and SSC transgenic cassava plants. Molecular mass standards are indicated on the left. The antibodies used for Western blotting are indicated on the bottom. Sample names are indicated on the top. Arrowheads indicate the bands that match the predicted molecular weights.

Furthermore, to investigate the effect of MeSSII on starch granule-bound protein, granule-bound proteins isolated from starch samples extracted from storage roots of WT and the SSC lines were detected by Western blotting (Fig. 6B). MeSSI, MeSBEII, MeISAI, and MeISAII were not detected in the SSC lines, although they were present as granule-bound proteins in WT, whereas no differences were observed in the granule-bound protein profiles for MeSBEI and MeGBSSI between WT and the SSC lines.

## Discussion

At present, all members of the starch synthase family in plants are classified into seven subfamilies that include GBSS, SSI, SSII, SSIII, SSIV, SSV, and SSVI based on the earliest available genomic sequences (Yang, Wang et al. 2013, Helle, Bray et al. 2018, Abt, Pfister et al. 2020), and mainly limited to Arabidopsis and economically important crop species. The functions of six starch synthases, including GBSSI, SSI, SSII, SSIII, SSIV, and SSV have been well-documented in previous studies (Craig, Lloyd et al. 1998, Edwards, Fulton et al. 1999, Kossmann, Abel et al. 1999, McPherson and Jane 1999, Song and Jane 2000, Yamamori, Fujita et al. 2000, Fulton, Edwards et al. 2002, Yoo and Jane 2002, Morell, Kosar-Hashemi et al. 2003, Delvalle, Dumez et al. 2005, Nakamura, Francisco et al. 2005, Fujita, Yoshida et al. 2006, Fujita, Yoshida et al. 2007, Roldan, Wattebled et al. 2007, Ryoo, Yu et al. 2007, Zhang, Szydlowski et al. 2008, Takahata, Tanaka et al. 2010, Zhao, Dufour et al. 2011, Zhou, Wang et al. 2016, Wang, Li et al. 2017, Abt, Pfister et al. 2020). It has been demonstrated that amylopectin biosynthetic isozymes can interact with each other in the endosperms of wheat, maize, barley, and rice seeds (Tetlow, Wait et al. 2004, Hennen-Bierwagen, Liu et al. 2008, Tetlow, Beisel et al. 2008, Hennen-Bierwagen, Lin et al. 2009, Liu, Makhmoudova et al. 2009, Ahmed, Tetlow et al. 2015, Crofts, Abe et al. 2015). Also, amylose synthase GBSSI is also able to form oligomers in rice, and GBSSI activity and oligomerization are affected by phosphorylation, redox regulation, and ADPG (Liu, Huang et al. 2013). Nevertheless, these knowledges are limited to the seeds of the major gramineous crop species. It is still unclear whether similar starch synthase complexes are present in dicotyledonous species and in underground storage organs. In this study, by focusing on cassava SSII, we have elucidated its function and the molecular mechanism by which it chaperones the starch synthase complex to regulate starch biosynthesis in cassava storage roots (Figs. 2-6, Supplementary Fig. S1-2 and Table S2-3). Based on our findings and those of previously published studies (Hennen-Bierwagen, Lin et al. 2009, Crofts, Abe et al. 2015), we propose a functional model of MeSSII that uncovers new features of starch biosynthesis in dicot plants, especially in root crops (Fig. 7).

**Figure 7.**
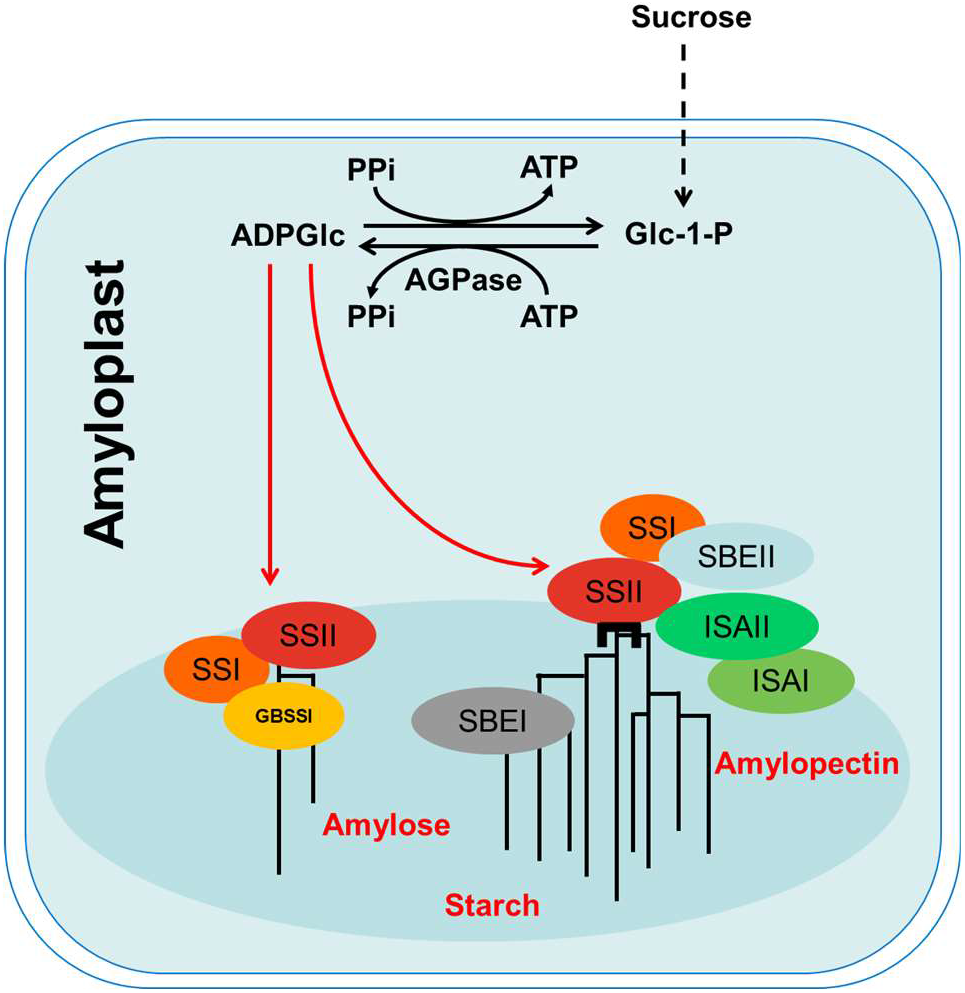
A molecular model of SSII function in starch biosynthesis in cassava storage roots. SSII influences the assembly of heteromeric starch synthetic protein complexes and the starch binding ability of SSI, SBEII, ISAI, and ISAII through protein-protein interactions. Down-regulated *SSII* expression disrupts these protein complexes, which results in pronounced changes of starch properties in cassava storage roots.

Further investigation showed that MeSSII is important for growth and development in cassava (Fig. 2D and Supplementary Fig. S1), and the starch isolated from storage roots of the *MeSSII*-RNAi lines showed altered physico-chemical properties, such as amylose content, pasting, gelatinization properties, and crystallinity (Fig. 3 and Supplementary Fig. S2 and Table S2-3), indicating that MeSSII plays a vital role in starch biosynthesis in cassava. Among these properties, the increased amylose content in roots of the SSC lines is an interesting phenomenon (Fig. 3C), and implies that as an amylopectin synthase, MeSSII has an important influence on amylose synthesis. Analyses of the mRNA and protein expression levels of key enzymes involved in starch synthesis demonstrated that the effect of MeSSII on starch biosynthesis was not achieved by affecting the expression levels of the genes that encode these enzymes (Supplementary Fig. S5A, B). Our results show that MeSSII influenced the ability of MeSSI, MeSBEII, MeISAI, and MeISAII to bind to starch granules (Fig. 6B), which is similar to the effect of cereal SSIIa on the ability of SSI and SBEII to bind to starch granule (Yamamori, Fujita et al. 2000, Morell, Kosar-Hashemi et al. 2003, Umemoto and Aoki 2005), although the presence of a carbohydrate binding module was not predicted in MeSSII (Fig. 1B). This indicates that MeSSII may bind to starch granules by interacting with other proteins that contain a CBM domain. Furthermore, obvious changes in the GPC mobility of MeSSI and MeSBEII in the *MeSSII*-RNAi lines also implies the significance of MeSSII in these protein complexes that consist of starch biosynthetic enzymes (Fig. 6A). Therefore, we propose that owing to the disruption of these protein complexes, a deficiency in MeSSII affects the starch binding ability of other key enzymes in starch synthesis and even might influence the activity of these enzymes, leading to a significant alteration in starch properties.

In this study, we have confirmed the presence of multienzyme complexes for starch biosynthesis in cassava storage roots, a dicot species. The results of our study also revealed previously-unknown protein interactions among starch biosynthetic enzymes in plants. Although protein-protein interactions among SSs including GBSSI have been reported (Liu, Huang et al. 2013, Crofts, Abe et al. 2015, Seung, Soyk et al. 2015), it is still unclear whether GBSSI participates in amylopectin synthesis. Here, we found that MeGBSSI can interact with MeSSI and MeISAI directly (Fig. 5A, B), strongly implying that GBSSI is directly involved in the process of amylopectin synthesis. With regard to the function of the interactions between MeGBSSI and MeSSI or MeISAI, GBSSI may interact with amylopectin enzymatic complexes through SSI, and contribute to the synthesis of extra-long amylopectin chains directly, as shown in previous studies in potato and rice (Fulton, Edwards et al. 2002, Hanashiro, Itoh et al. 2008). Furthermore, GBSSI may interact with SBEII through SSI to produce more branches in amylose and the extra-long amylopectin chains, leading to the synthesis of more amylopectin clusters. It should also be noted that pea amylose was reported to be debranched completely by isoamylase (ISA) in aqueous 40% dimethyl sulfoxide (Colonna and Mercier 1984). The interaction between GBSSI and ISAI indicates that ISAI may excise the extra erroneous branched chains in amylopectin in favor of producing long chains or the synthesis of amylose by interacting with GBSSI. In *Chlamydomonas reinhardtii*, amylose can be formed by extending the branches on amylopectin, which are then removed from the amylopectin chains (van de Wal, D’Hulst et al. 1998). The interaction between GBSSI and the amylopectin enzymatic complexes through SSI might play a role in regulating the activity of one another. In wheat, SSI, SSII, SBEIIa, or SBEIIb can form heterotrimers after phosphorylation of SBEIIa or SBEIIb, which increases the enzymatic activity of SBEIIa and SBEIIb significantly (Tetlow, Beisel et al. 2008). In addition, PhoI can improve the activity of SBE by interacting with SBE in rice (Crofts, Abe et al. 2015).

To date, a direct interaction between ISAs and the protein complexes that consist of SS and SBE has not been reported. The removal of erroneous branches by ISAs plays an important role in the extension and branching of amylopectin chains, and, therefore, protein interactions with ISAs are essential (Hussain, Mant et al. 2003, Delatte, Trevisan et al. 2005, Kubo, Colleoni et al. 2010, Utsumi, Utsumi et al. 2011, Peng, Wang et al. 2014). Lin *et al*. (2012) hypothesized that ISAI may interact with SSIII, either directly or indirectly, so that ISAI can bind to the growing amylopectin molecules (Lin, Huang et al. 2012). The interaction between ISA and a CBM-containing FLO6 protein shows that ISA could potentially bind to starch granules via FLO6 (Peng, Wang et al. 2014). In cassava, we found that MeSSII can interact with MeISAII directly (Fig. 5A, B), suggesting that MeISAI and MeISAII are part of the protein complexes that consist of SS and SBE via their interactions with MeSSII, and hence, contributing directly to amylopectin synthesis by removing the erroneous branches.

In conclusion, we elucidated the function of MeSSII in starch biosynthesis in cassava storage roots. Our work describes a new mechanism of starch accumulation in underground storage organs of a dicot species. Contrary to current dogma, our results suggest that starch synthesis requires a close coordination between GBSSI and amylopectin biosynthetic enzymes, and that amylose synthesis and amylopectin synthesis are directly interrelated. Further molecular investigations into the regulatory mechanisms of starch biosynthesis in cassava, such as the transcription factors that regulate the expression of the genes that encode these enzymes, will improve our understanding of the mechanisms governing starch accumulation in cassava storage roots. Additionally, this work provides a novel source of cassava starch with a high amylose content for various industrial applications.

## Materials and methods

### Phylogenetic tree analysis of starch synthases

Arabidopsis (*Arabidopsis thaliana*) SSI, SSII, SSIII, SSIV, and SSV sequences were retrieved from Phytozome and the closest orthologs in the other species, including *Amborella trichopoda, Brachypodium distachyon, Capsella rubella, Chlamydomonas reinhardtii, Glycine max, Gossypium raimondii, Hordeum vulgare, Manihot esculenta, Oryza sativa, Populus trichocarpa, Physcomitrella patens, Prunus persica, Sorghum bicolor, Vitis vinifera* and *Zea mays*, were obtained from the Phytozome and Ensembl Plants via BLASTp. The three closest prokaryotic glycogen synthase sequences of AtSSIII, At-SSIV, and At-SSV (WP 048672006.1, WP 059061331.1, and WP026046064.1, respectively) were retrieved from the National Center for Biotechnology Information as an outgroup. We aligned sequences using MUSCLE (Edgar 2004) with the default parameters that are tuned for accuracy. A phylogenetic tree was created by MEGA7, using the maximum likelihood method with the Jones-Taylor-Thornton matrix-based model (Jones, Taylor et al. 1992). Bootstrap values were obtained with 1,000 replications. Positions with less than 95% site coverage were eliminated. The exon-intron structure and conserved amino acid domains of these SSII genes was determined by the Gene Structure Display Server (GSDS: http://gsds.gao-lab.org/index.php).

### *MeSSII* cloning and production of *MeSSII-*RNAi transgenic cassava

Sequence information for *MeSSII* was derived from the cassava genome database (http://www.phytozome.net). The full length cDNA of *MeSSII* was cloned from a cassava cDNA library. The binary RNAi gene silencing expression vector p35S::*MeSSII*-RNAi was constructed from the plasmid pRNAi-dsAC1 (Vanderschuren, Alder et al. 2009). The AC1 sequence was excised and replaced with a partial *MeSSII* cDNA sequence from 978 bp to 1177 bp. The control construct was pRNAi-dsAC1 without the *MeSSII* cDNA. The two constructs were introduced into *Agrobacterium tumefaciens* LBA4404 and the cassava cultivar TMS60444 was used to generate transgenic plants as described previously by Zhang *et al*. (2000) (Zhang, Potrykus et al. 2000). The integration pattern of T-DNA in transgenic cassava was determined by Southern blot analysis as described by Zhao *et al*. (2011) (Zhao, Dufour et al. 2011). The DIG-labeled hygromycin phosphotransferase (HPT) gene was used as the hybridization probe.

The regenerated transgenic lines and WT plants were propagated *in vitro* and then transferred to pots in the greenhouse for macro-propagation (16 h/8 h of light/dark, 30°C/22°C day/night). The *MeSSII*-RNAi transgenic lines are given the designation SSC, which is an abbreviation for the “starch synthase catalytic domain” used for constructing the p35S::*MeSSII*-RNAi vector, and the control lines are designated EV, which is the abbreviation for “empty vector”. For field evaluation, 10 stems per transgenic line and WT were planted in early May in the Wushe Plantation for Transgenic Crops, Shanghai, China (31°13948.0099 N, 121°28912.0099E), and harvested in early November. The phenotypic performance of the plants was recorded.

### Transcriptional and translational expression analysis

Leaves, petioles, stems, and storage roots from at least three plants per line were harvested and then ground to a fine powder in liquid N_2_ for mRNA extraction. To quantify the expression of genes, qRT-PCR was performed as described (Xu, Duan et al. 2013). The *β-Actin* gene was used as a reference to normalize gene expression. The sequences of the PCR primers used in these experiments are shown in Table S5.

Protein extraction was performed as described by Crofts *et al*. (2015) (Crofts, Abe et al. 2015) with some modifications. Mature cassava storage roots (diameter >2.5 cm; 1 g samples) were used for each extraction. Total protein was extracted with 1 vol (w/v) of a denaturing buffer containing 0.125 M Tris-HCl (pH 6.8), 8 M urea, 4% SDS, and 5% β-mercaptoethanol. Samples were extracted overnight at room temperature, centrifuged at 12,000 rpm to remove gelatinized starch and other particulate matter, and the supernatants were used for denaturing polyacrylamide gel electrophoresis (SDS-PAGE) and immunoblotting. Soluble proteins were extracted on ice with 6 vols (w/v) (three repeats with 2 vols each) of an extraction buffer containing 10 mM HEPES-KOH (pH 7.5) and 100 mM NaCl. After extraction, samples were centrifuged at 12,000 rpm at 4°C for 15 min. The residual pellet was extracted with 1 vol (w/v) of denaturing buffer as described above, centrifuged as before, and the supernatant was then used to represent the insoluble, starch granule-associated proteins.

Proteins were transferred to polyvinylidene fluoride (PVDF) membranes after SDS-PAGE or blue native (BN)-PAGE. Membranes were treated as described by Crofts *et al*. (2015) (Crofts, Abe et al. 2015). Polyclonal antibodies against MeSSI, MeSSII, MeSBEI, MeSBEII, MeISAI, MeISAII, and MeGBSSI were prepared by ABclonal Biotech Co., Ltd using purified recombinant proteins (purity >95%) as antigens. Primary antibodies were used at a dilution of 1:1,000. HRP-conjugated goat anti-rabbit IgG (Sigma) was used as a secondary antibody at a dilution of 1:10,000. The colorimetric signal was detected with the Pierce ECL Western Blotting Substrate Reagent Kit (catalog no. 32106; Thermo Fisher) and was visualized using the Tanon-5500 Chemiluminescent Imaging System (Tanon Science and Technology, Shanghai, China).

### Physico-chemical property assays of cassava storage starches

The granule size distribution of starch extracted from cassava storage roots was determined as described by Zhou *et al*. (2015) (Zhou, Yang et al. 2015). A Mastersizer 2000 laser diffraction instrument (Malvern Instruments Ltd., Worcestershire, UK) was used in wet-well mode. The starch was added to the reservoir and sonicated for 30 s at 6 W until an obscuration value of 12-17% was reached. The refractive indices used for the water and starch were 1.330 and 1.50, respectively. The particle size distributions are presented as the diameter versus volume. To examine whether starch granule morphology was altered in the different transgenic lines, the starch granules were coated with gold after they were spread on a metal stub and then observed under a scanning electron microscope (SEM, JSM6360lv, JEOL, Tokyo, Japan). Pieces of fresh mature roots (2 mm^3^) were coated with 1,3-diformal propane and subjected to ultra-thin sectioning. The sections were then photographed under a transmission electron microscope (TEM; Hitachi H7650, Tokyo, Japan).

The chain length distribution (CLD) of the starches was measured as described by Zhou *et al*. (2015) (Zhou, Yang et al. 2015). Starch from the transgenic plants and WT was digested with *Pseudomonas amyloderamosa* isoamylase (Sigma), and the CLD of the amylopectin was analyzed using high-performance anion-exchange chromatography with pulsed amperometric detection (HPAEC-PAD; Dionex-ICS 3000; Dionex Corporation, USA). A D8 Advance Bruker X-ray diffractometer (Bruker AXS, Germany) was used to study the crystal type of the starch granules. Starch powders were scanned with the 2*θ* diffraction angle at 4-50°. The crystal patterns and degree of crystallinity were analyzed using Jade 5.0 software.

The thermal properties of the starch samples were determined using a differential scanning calorimeter (DSC Q2000; TA Instrument Ltd., UK). A mixture of starch and distilled water (w/w, 1:3) was sealed in an aluminum pan and scanned at 10°C/min from 30°C to 95°C with an empty aluminum pan as a reference. The gelatinization temperature and enthalpy were calculated using Universal Analysis 2000 software. The pasting properties of the starch were analyzed using a rapid viscosity analyzer (model RVA-Super 3; Newport Scientific Pty. Ltd., Australia). The starch was suspended in distilled water (5% w/v, dry weight basis) and tested using a dedicated program for cassava. The temperature of the starch slurry was increased from 30°C to 95°C at a rate of 5°C/min and held at 95°C for 6 min, followed by cooling to 50°C at the same rate, and the temperature was maintained at 50°C for 10 min. The rotating speed of the paddle was 960 rpm during the first 10 s, after which it was kept constant (160 rpm) throughout the analysis.

The amylose content of the starch was measured using colorimetric amylose content determination as previously described (Knutson and Grove 1994). Amylose (Type III, Sigma A0512, St. Louis, MO) and amylopectin (Sigma 10118) from potato were used to establish standard curves.

### Analysis of protein interactions

Samples of cassava storage roots (1.5 g) were extracted in 1.5 ml of gel filtration buffer containing 50 mM Tris-acetate (pH 7.5), 100 mM NaCl, 10 μl/ml plant protease inhibitor cocktail (Sigma-Aldrich LLC) and centrifuged at 12,000 rpm for 15 min. The supernatants were injected into a 500 μl sample loop, and fractionated by gel permeation chromatography (GPC) using a Superdex 200 10/300 GL column (GE Healthcare, catalog no. 17-5175-01) connected to an AKTA FPLC system (GE Healthcare) at 4°C. The column was equilibrated with 50 mM Tris-acetate buffer (pH 7.5) containing 100 mM NaCl; the flow rate was 1 ml/min and fraction sizes of 0.5 ml were collected. Molecular mass standards run under identical conditions were from GE Healthcare (catalog no. 17-0445-01). Samples from each fraction (40 μl) were analyzed by SDS-PAGE and immunoblotting.

The BN-PAGE procedures used in this study were performed as previously described (Crofts et al., 2015) (Crofts, Abe et al. 2015) with some modifications. Cassava storage roots were extracted with 1 vol (w/v) of 50 mM Bis-Tris (pH 7.5), 50 mM NaCl, 10% glycerol, 0.001% Ponceau S, and centrifuged at 12,000 × *g* for 10 min. The supernatants were subjected to 3-12% acrylamide Bis-Tris native-PAGE (Invitrogen, catalog no. BN1001BOX) and electrophoresed with an anode buffer containing 50 mM Bis-Tris and 50 mM tricine, and a cathode buffer containing 50 mM Bis-Tris, 50 mM tricine and 0.004% CBB G-250 stain at 80 V for an initial 1 h and at 120 V for the remaining time.

Co-immunoprecipitation experiments were performed using the methods described by Liu *et al*. (2012) (Liu, Ahmed et al. 2012). Purified peptide-specific antibodies (each approximately 2 μg) were individually used for the co-immunoprecipitation (Co-IP) experiments with the soluble protein extracts described above (1 ml, between 0.8 and 1.2 mg/ml protein). The protein extract/antibody mixtures were incubated at room temperature on a rotator for 50 min, and immunoprecipitation of the antibody/protein complexes was performed by adding 50 μl of Protein A-Sepharose (Sigma-Aldrich) made up as a 50% (w/v) slurry with phosphate buffered saline (PBS; 137 mM NaCl, 10 mM Na_2_HPO_4_, 2.7 mM KCl, 1.8 M KH_2_PO_4_, pH 7.4) at room temperature for 40 min. The Protein A-Sepharose/antibody/protein complexes were recovered by centrifugation at 2,000 × *g* for 5 min at 4°C in a refrigerated microcentrifuge, and the supernatants were discarded. The pellets were washed five times with PBS (1.3 ml each), followed by washing five times with a buffer containing 10 mM HEPES-NaOH (pH 7.5) and 150 mM NaCl. The washed pellets were boiled in SDS-containing gel loading buffer and separated by SDS-PAGE, followed by immunoblot analysis. In order to exclude the possibility that co-immunoprecipitation of the proteins observed in the immunoprecipitation pellet was a result of AGPL, AGPS, SS, SBE, ISA or GBSS binding to the same glucan chain, the soluble protein extracts used for immunoprecipitation were pre-incubated with 5 U each of the glucan degrading enzymes amyloglucosidase (EC 3.2.1.3, Sigma A7255) and α-amylase (EC 3.2.1.1, Sigma A2643) for 20 min at 25°C.

Yeast two-hybridization (Y2H) assays were performed according to the manufacture’s procedure (Clontech Laboratories Inc., Palo Alto, CA, USA). For each isozyme, a DNA fragment that was PCR-amplified using the specific primer pair (Table S1) was cloned in-frame into pGBKT7 to form the bait construct. The same fragment was inserted into pGADT7 to form the prey construct. Both plasmids were transformed into yeast cells (strain AH109). The resulting yeast transformants were grown on SD medium containing 3-AT and X-α-gal, but lacking Trp, Leu, His, and Ade, for 2-3 days at 30°C.

Bimolecular fluorescence complementation was conducted as previously described (Ma, Kong et al. 2015) with some modifications. Each starch biosynthetic enzyme cDNA was introduced into BiFC vectors enabling protein fusions to the N-terminal half of YFP or C-terminal half of YFP (nYFP-fused and cYFP-fused, respectively). nYFP-fused with cYFP-fused, nYFP-fused with cYFP, nYFP with cYFP-fused, or nYFP-cYFP cDNAs were transiently co-produced in *N. benthamiana* leaves. Confocal images of *N. benthamiana* epidermal cells transiently co-expressing these cDNAs are used to analyze these interactions here. The primers used in these experiments are shown in Table S5.

### Mass spectrometry

The proteins were digested in the gel with bovine trypsin as previously described (Wang, Li et al. 2007). After digestion, the peptides were collected and detected by LC-MS/MS according to the facility procedures used at the Public Mass Spectrometry Technical Service Center, CAS Center for Excellence in Molecular Plant Sciences, Chinese Academy of Sciences. Data were analyzed using Proteome Discoverer software.

### Statistical analysis

Root samples were collected from three independent plants per line. Data from at least three replicates are presented as mean ± SD. Analysis of independent samples with Student’s *t*-test was performed using SPSS software, version 17 (SPSS Inc., Chicago, IL, USA). An alpha value of *P* <0.05 was considered to be statistically significant.

## Acknowledgements

We thank Xinyan Liu and Chuanzhong Li for assistance in field experiments; Xiaoyan Gao, Zhiping Zhang, and Jiqin Li for assistance with the SEM and TEM experiments; and Yuanhong Shan for assistance with LC-MS/MS analysis. This work was supported by the National Science Foundation of China (No. 31871682), the National Key Research and Development Program of China (Nos. 2017YFA0505500, 2019YFD1001100), the Earmarked Fund for China Agriculture Research System (CARS-11-shzp), and Shanghai Municipal Science and Technology Major Project (No. 2017SHZDZX01).

## Author contributions

S.H. performed most of the experiments, analyzed the data, and drafted the manuscript; X.H., S.W., and W.Z. conducted some of the experiments; S.W. and Q.M. produced the transgenic cassava plants; X.H., Q.M. and X.L. maintained the transgenic cassava plants; P.Z., L.C., S.H. designed and conceived this project; P.Z. and L.C. analyzed the data and revised the manuscript with input from the other authors. All authors contributed to and approved the final manuscript.

## References

Abe, N., H. Asai, H. Yago, N. F. Oitome, R. Itoh, N. Crofts, Y. Nakamura and N. Fujita (2014). “Relationships between starch synthase I and branching enzyme isozymes determined using double mutant rice lines.” Bmc Plant Biology 14.

Abt, M. R., B. Pfister, M. Sharma, S. Eicke, L. Burgy, I. Neale, D. Seung and S. C. Zeeman (2020). “STARCH SYNTHASE5, a Noncanonical Starch Synthase-Like Protein, Promotes Starch Granule Initiation in Arabidopsis.” Plant Cell 32(8): 2543–2565.

Ahmed, Z., I. J. Tetlow, R. Ahmed, M. K. Morell and M. J. Emes (2015). “Protein-protein interactions among enzymes of starch biosynthesis in high-amylose barley genotypes reveal differential roles of heteromeric enzyme complexes in the synthesis of A and B granules.” Plant Sci 233: 95–106.

Bull, S. E., D. Seung, C. Chanez, D. Mehta, J. E. Kuon, E. Truernit, A. Hochmuth, I. Zurkirchen, S. C. Zeeman, W. Gruissem and H. Vanderschuren (2018). “Accelerated ex situ breeding of GBSS-and PTST1-edited cassava for modified starch.” Sci Adv 4(9): eaat6086.

Ceballos, H., T. Sanchez, N. Morante, M. Fregene, D. Dufour, A. M. Smith, K. Denyer, J. C. Perez, F. Calle and C. Mestres (2007). “Discovery of an amylose-free starch mutant in cassava (Manihot esculenta Crantz).” J Agric Food Chem 55(18): 7469–7476.

Cock, J. H. (1982). “Cassava: a basic energy source in the tropics.” Science 218(4574): 755–762.

Colonna, P. and C. Mercier (1984). “Macromolecular Structure of Wrinkled-Pea and Smooth-Pea Starch Components.” Abstracts of Papers of the American Chemical Society 188(Aug): 59-Carb.

Craig, J., J. R. Lloyd, K. Tomlinson, L. Barber, A. Edwards, T. L. Wang, C. Martin, C. L. Hedley and A. M. Smith (1998). “Mutations in the gene encoding starch synthase II profoundly alter amylopectin structure in pea embryos.” Plant Cell 10(3): 413–426.

Crofts, N., N. Abe, N. F. Oitome, R. Matsushima, M. Hayashi, I. J. Tetlow, M. J. Emes, Y. Nakamura and N. Fujita (2015). “Amylopectin biosynthetic enzymes from developing rice seed form enzymatically active protein complexes.” J Exp Bot 66(15): 4469–4482.

Delatte, T., M. Trevisan, M. L. Parker and S. C. Zeeman (2005). “Arabidopsis mutants Atisa1 and Atisa2 have identical phenotypes and lack the same multimeric isoamylase, which influences the branch point distribution of amylopectin during starch synthesis.” Plant Journal 41(6): 815–830.

Delvalle, D., S. Dumez, F. Wattebled, I. Roldan, V. Planchot, P. Berbezy, P. Colonna, D. Vyas, M. Chatterjee, S. Ball, A. Merida and C. D’Hulst (2005). “Soluble starch synthase I: a major determinant for the synthesis of amylopectin in Arabidopsis thaliana leaves.” Plant J 43(3): 398–412.

Edgar, R. C. (2004). “MUSCLE: multiple sequence alignment with high accuracy and high throughput.” Nucleic Acids Res 32(5): 1792–1797.

Edwards, A., D. C. Fulton, C. M. Hylton, S. A. Jobling, M. Gidley, U. Rossner, C. Martin and A. M. Smith (1999). “A combined reduction in activity of starch synthases II and III of potato has novel effects on the starch of tubers.” Plant Journal 17(3): 251–261.

Fujita, N., M. Yoshida, N. Asakura, T. Ohdan, A. Miyao, H. Hirochika and Y. Nakamura (2006). “Function and characterization of starch synthase I using mutants in rice.” Plant Physiology 140(3): 1070–1084.

Fujita, N., M. Yoshida, T. Kondo, K. Saito, Y. Utsumi, T. Tokunaga, A. Nishi, H. Satoh, J. H. Park, J. L. Jane, A. Miyao, H. Hirochika and Y. Nakamura (2007). “Characterization of SSIIIa-deficient mutants of rice: the function of SSIIIa and pleiotropic effects by SSIIIa deficiency in the rice endosperm.” Plant Physiol 144(4): 2009–2023.

Fulton, D. C., A. Edwards, E. Pilling, H. L. Robinson, B. Fahy, R. Seale, L. Kato, A. M. Donald, P. Geigenberger, C. Martin and A. M. Smith (2002). “Role of granule-bound starch synthase in determination of amylopectin structure and starch granule morphology in potato.” J Biol Chem 277(13): 10834–10841.

Grimaud, F., H. Rogniaux, M. G. James, A. M. Myers and V. Planchot (2008). “Proteome and phosphoproteome analysis of starch granule-associated proteins from normal maize and mutants affected in starch biosynthesis.” J Exp Bot 59(12): 3395–3406.

Hanashiro, I., K. Itoh, Y. Kuratomi, M. Yamazaki, T. Igarashi, J. I. Matsugasako and Y. Takeda (2008). “Granule-bound starch synthase I is responsible for biosynthesis of extra-long unit chains of amylopectin in rice.” Plant and Cell Physiology 49(6): 925–933.

Helle, S., F. Bray, J. Verbeke, S. Devassine, A. Courseaux, M. Facon, C. Tokarski, C. Rolando and N. Szydlowski (2018). “Proteome Analysis of Potato Starch Reveals the Presence of New Starch Metabolic Proteins as Well as Multiple Protease Inhibitors.” Frontiers in Plant Science 9.

Hennen-Bierwagen, T. A., Q. Lin, F. Grimaud, V. Planchot, P. L. Keeling, M. G. James and A. M. Myers (2009). “Proteins from multiple metabolic pathways associate with starch biosynthetic enzymes in high molecular weight complexes: a model for regulation of carbon allocation in maize amyloplasts.” Plant Physiol 149(3): 1541–1559.

Hennen-Bierwagen, T. A., F. Liu, R. S. Marsh, S. Kim, Q. Gan, I. J. Tetlow, M. J. Emes, M. G. James and A. M. Myers (2008). “Starch biosynthetic enzymes from developing maize endosperm associate in multisubunit complexes.” Plant Physiol 146(4): 1892–1908.

Hizukuri, S. (1985). “Relationship between the Distribution of the Chain-Length of Amylopectin and the Crystalline-Structure of Starch Granules.” Carbohydrate Research 141(2): 295–306.

Hizukuri, S. (1986). “Polymodal Distribution of the Chain Lengths of Amylopectins, and Its Significance.” Carbohydrate Research 147(2): 342–347.

Hussain, H., A. Mant, R. Seale, S. Zeeman, E. Hinchliffe, A. Edwards, C. Hylton, S. Bornemann, A. M. Smith, C. Martin and R. Bustos (2003). “Three isoforms of isoamylase contribute different catalytic properties for the debranching of potato glucans.” Plant Cell 15(1): 133–149.

Jeon, J. S., N. Ryoo, T. R. Hahn, H. Walia and Y. Nakamura (2010). “Starch biosynthesis in cereal endosperm.” Plant Physiol Biochem 48(6): 383–392.

Jones, D. T., W. R. Taylor and J. M. Thornton (1992). “The rapid generation of mutation data matrices from protein sequences.” Comput Appl Biosci 8(3): 275–282.

Knutson, C. A. and M. J. Grove (1994). “Rapid Method for Estimation of Amylose in Maize Starches.” Cereal Chemistry 71(5): 469–471.

Kossmann, J., G. J. W. Abel, F. Springer, J. R. Lloyd and L. Willmitzer (1999). “Cloning and functional analysis of a cDNA encoding a starch synthase from potato (Solanum tuberosum L,) that is predominantly expressed in leaf tissue.” Planta 208(4): 503–511.

Kubo, A., C. Colleoni, J. R. Dinges, Q. Lin, R. R. Lappe, J. G. Rivenbark, A. J. Meyer, S. G. Ball, M. G. James, T. A. Hennen-Bierwagen and A. M. Myers (2010). “Functions of heteromeric and homomeric isoamylase-type starch-debranching enzymes in developing maize endosperm.” Plant Physiol 153(3): 956–969.

Li, S., Y. Cui, Y. Zhou, Z. Luo, J. Liu and M. Zhao (2017). “The industrial applications of cassava: current status, opportunities and prospects.” J Sci Food Agric 97(8): 2282–2290.

Lin, Q. H., B. Q. Huang, M. X. Zhang, X. L. Zhang, J. Rivenbark, R. L. Lappe, M. G. James, A. M. Myers and T. A. Hennen-Bierwagen (2012). “Functional Interactions between Starch Synthase III and Isoamylase-Type Starch-Debranching Enzyme in Maize Endosperm.” Plant Physiology 158(2): 679–692.

Liu, D. R., W. X. Huang and X. L. Cai (2013). “Oligomerization of rice granule-bound starch synthase 1 modulates its activity regulation.” Plant Sci 210: 141–150.

Liu, F., Z. Ahmed, E. A. Lee, E. Donner, Q. Liu, R. Ahmed, M. K. Morell, M. J. Emes and I. J. Tetlow (2012). “Allelic variants of the amylose extender mutation of maize demonstrate phenotypic variation in starch structure resulting from modified protein-protein interactions.” J Exp Bot 63(3): 1167–1183.

Liu, F., N. Romanova, E. A. Lee, R. Ahmed, M. Evans, E. P. Gilbert, M. K. Morell, M. J. Emes and I. J. Tetlow (2012). “Glucan affinity of starch synthase IIa determines binding of starch synthase I and starch-branching enzyme IIb to starch granules.” Biochem J 448(3): 373–387.

Liu, F. S., A. Makhmoudova, E. A. Lee, R. Wait, M. J. Emes and I. J. Tetlow (2009). “The amylose extender mutant of maize conditions novel protein-protein interactions between starch biosynthetic enzymes in amyloplasts.” Journal of Experimental Botany 60(15): 4423–4440.

Liu, H., G. Yu, B. Wei, Y. Wang, J. Zhang, Y. Hu, Y. Liu, G. Yu, H. Zhang and Y. Huang (2015). “Identification and Phylogenetic Analysis of a Novel Starch Synthase in Maize.” Front Plant Sci 6: 1013.

Liu, K., Y. Y. Zu, C. D. Chi, B. Gu, L. Chen and X. X. Li (2018). “Modulation of the digestibility and multi-scale structure of cassava starch by controlling the cassava growth period.” International Journal of Biological Macromolecules 120: 346–353.

Ma, W., Q. Kong, M. Grix, J. J. Mantyla, Y. Yang, C. Benning and J. B. Ohlrogge (2015). “Deletion of a C-terminal intrinsically disordered region of WRINKLED1 affects its stability and enhances oil accumulation in Arabidopsis.” Plant J 83(5): 864–874.

Makhmoudova, A., D. Williams, D. Brewer, S. Massey, J. Patterson, A. Silva, K. A. Vassall, F. Liu, S. Subedi, G. Harauz, K. W. Siu, I. J. Tetlow and M. J. Emes (2014). “Identification of multiple phosphorylation sites on maize endosperm starch branching enzyme IIb, a key enzyme in amylopectin biosynthesis.” J Biol Chem 289(13): 9233–9246.

McPherson, A. E. and J. Jane (1999). “Comparison of waxy potato with other root and tuber starches.” Carbohydrate Polymers 40(1): 57–70.

Mehrpouyan, S., U. Menon, I. J. Tetlow and M. J. Emes (2020). “Protein Phosphorylation Regulates Maize Endosperm Starch Synthase IIa Activity and Protein-Protein Interactions.” Plant J.

Moorthy, S. N. (2002). “Physicochemical and functional properties of tropical tuber starches: A review.” Starch-Starke 54(12): 559–592.

Morell, M. K., B. Kosar-Hashemi, M. Cmiel, M. S. Samuel, P. Chandler, S. Rahman, A. Buleon, I. L. Batey and Z. Y. Li (2003). “Barley sex6 mutants lack starch synthase IIa activity and contain a starch with novel properties.” Plant Journal 34(2): 172–184.

Nakamura, Y. (2002). “Towards a better understanding of the metabolic system for amylopectin biosynthesis in plants: Rice endosperm as a model tissue.” Plant and Cell Physiology 43(7): 718–725.

Nakamura, Y., S. Aihara, N. Crofts, T. Sawada and N. Fujita (2014). “In vitro studies of enzymatic properties of starch synthases and interactions between starch synthase I and starch branching enzymes from rice.” Plant Science 224: 1–8.

Nakamura, Y., P. B. Francisco, Y. Hosaka, A. Sato, T. Sawada, A. Kubo and N. Fujita (2005). “Essential amino acids of starch synthase IIa differentiate amylopectin structure and starch quality between japonica and indica rice varieties.” Plant Molecular Biology 58(2): 213–227.

Nakamura, Y., M. Ono, C. Utsumi and M. Steup (2012). “Functional Interaction Between Plastidial Starch Phosphorylase and Starch Branching Enzymes from Rice During the Synthesis of Branched Maltodextrins.” Plant and Cell Physiology 53(5): 869–878.

Peng, C., Y. Wang, F. Liu, Y. Ren, K. Zhou, J. Lv, M. Zheng, S. Zhao, L. Zhang, C. Wang, L. Jiang, X. Zhang, X. Guo, Y. Bao and J. Wan (2014). “FLOURY ENDOSPERM6 encodes a CBM48 domain-containing protein involved in compound granule formation and starch synthesis in rice endosperm.” Plant J 77(6): 917–930.

Roldan, I., F. Wattebled, M. Mercedes Lucas, D. Delvalle, V. Planchot, S. Jimenez, R. Perez, S. Ball, C. D’Hulst and A. Merida (2007). “The phenotype of soluble starch synthase IV defective mutants of Arabidopsis thaliana suggests a novel function of elongation enzymes in the control of starch granule formation.” Plant J 49(3): 492–504.

Ryoo, N., C. Yu, C. S. Park, M. Y. Baik, I. M. Park, M. H. Cho, S. H. Bhoo, G. An, T. R. Hahn and J. S. Jeon (2007). “Knockout of a starch synthase gene OsSSIIIa/Flo5 causes white-core floury endosperm in rice (Oryza sativa L.).” Plant Cell Rep 26(7): 1083–1095.

Satoh, H., K. Shibahara, T. Tokunaga, A. Nishi, M. Tasaki, S. K. Hwang, T. W. Okita, N. Kaneko, N. Fujita, M. Yoshida, Y. Hosaka, A. Sato, Y. Utsumi, T. Ohdan and Y. Nakamura (2008). “Mutation of the plastidial alpha-glucan phosphorylase gene in rice affects the synthesis and structure of starch in the endosperm.” Plant Cell 20(7): 1833–1849.

Seung, D., J. Boudet, J. Monroe, T. B. Schreier, L. C. David, M. Abt, K. J. Lu, M. Zanella and S. C. Zeemana (2017). “Homologs of PROTEIN TARGETING TO STARCH Control Starch Granule Initiation in Arabidopsis Leaves.” Plant Cell 29(7): 1657-+.

Seung, D., S. Soyk, M. Coiro, B. A. Maier, S. Eicke and S. C. Zeeman (2015). “PROTEIN TARGETING TO STARCH Is Required for Localising GRANULE-BOUND STARCH SYNTHASE to Starch Granules and for Normal Amylose Synthesis in Arabidopsis.” Plos Biology 13(2).

Song, Y. and J. Jane (2000). “Characterization of barley starches of waxy, normal, and high amylose varieties.” Carbohydrate Polymers 41(4): 365–377.

Subasinghe, R. M., F. Liu, U. C. Polack, E. A. Lee, M. J. Emes and I. J. Tetlow (2014). “Multimeric states of starch phosphorylase determine protein-protein interactions with starch biosynthetic enzymes in amyloplasts.” Plant Physiol Biochem 83: 168–179.

Takahata, Y., M. Tanaka, M. Otani, K. Katayama, K. Kitahara, O. Nakayachi, H. Nakayama and M. Yoshinaga (2010). “Inhibition of the expression of the starch synthase II gene leads to lower pasting temperature in sweetpotato starch.” Plant Cell Rep 29(6): 535–543.

Tetlow, I. J., K. G. Beisel, S. Cameron, A. Makhmoudova, F. Liu, N. S. Bresolin, R. Wait, M. K. Morell and M. J. Emes (2008). “Analysis of protein complexes in wheat amyloplasts reveals functional interactions among starch biosynthetic enzymes.” Plant Physiology 146(4): 1878–1891.

Tetlow, I. J., R. Wait, Z. Lu, R. Akkasaeng, C. G. Bowsher, S. Esposito, B. Kosar-Hashemi, M. K. Morell and M. J. Emes (2004). “Protein phosphorylation in amyloplasts regulates starch branching enzyme activity and protein-protein interactions.” Plant Cell 16(3): 694–708.

Umemoto, T. and N. Aoki (2005). “Single-nucleotide polymorphisms in rice starch synthase IIa that alter starch gelatinisation and starch association of the enzyme.” Functional Plant Biology 32(9): 763–768.

Utsumi, Y., C. Utsumi, T. Sawada, N. Fujita and Y. Nakamura (2011). “Functional Diversity of Isoamylase Oligomers: The ISA1 Homo-Oligomer Is Essential for Amylopectin Biosynthesis in Rice Endosperm.” Plant Physiology 156(1): 61–77.

van de Wal, M., C. D’Hulst, J. P. Vincken, A. Buleon, R. Visser and S. Ball (1998). “Amylose is synthesized in vitro by extension of and cleavage from amylopectin.” Journal of Biological Chemistry 273(35): 22232–22240.

Vanderschuren, H., A. Alder, P. Zhang and W. Gruissem (2009). “Dose-dependent RNAi-mediated geminivirus resistance in the tropical root crop cassava.” Plant Mol Biol 70(3): 265–272.

Wang, X., X. Li, X. Deng, H. Han, W. Shi and Y. Li (2007). “A protein extraction method compatible with proteomic analysis for the euhalophyte Salicornia europaea.” Electrophoresis 28(21): 3976–3987.

Wang, Y., Y. Li, H. Zhang, H. Zhai, Q. Liu and S. He (2017). “A soluble starch synthase I gene, IbSSI, alters the content, composition, granule size and structure of starch in transgenic sweet potato.” Sci Rep 7(1): 2315.

Xu, J., X. Duan, J. Yang, J. R. Beeching and P. Zhang (2013). “Enhanced reactive oxygen species scavenging by overproduction of superoxide dismutase and catalase delays postharvest physiological deterioration of cassava storage roots.” Plant Physiol 161(3): 1517–1528.

Yamamori, M., S. Fujita, K. Hayakawa, J. Matsuki and T. Yasui (2000). “Genetic elimination of a starch granule protein, SGP-1, of wheat generates an altered starch with apparent high amylose.” Theoretical and Applied Genetics 101(1-2): 21–29.

Yang, Z., Y. Wang, S. Xu, C. Xu and C. Yan (2013). “Molecular evolution and functional divergence of soluble starch synthase genes in cassava (manihot esculenta crantz).” Evol Bioinform Online 9: 239–249.

Yoo, S. H. and J. L. Jane (2002). “Structural and physical characteristics of waxy and other wheat starches.” Carbohydrate Polymers 49(3): 297–305.

Zhang, P., I. Potrykus and J. Puonti-Kaerlas (2000). “Efficient production of transgenic cassava using negative and positive selection.” Transgenic Res 9(6): 405–415.

Zhang, X., N. Szydlowski, D. Delvalle, C. D’Hulst, M. G. James and A. M. Myers (2008). “Overlapping functions of the starch synthases SSII and SSIII in amylopectin biosynthesis in Arabidopsis.” BMC Plant Biol 8: 96.

Zhao, S. S., D. Dufour, T. Sanchez, H. Ceballos and P. Zhang (2011). “Development of waxy cassava with different Biological and physico-chemical characteristics of starches for industrial applications.” Biotechnol Bioeng 108(8): 1925–1935.

Zhou, H., L. Wang, G. Liu, X. Meng, Y. Jing, X. Shu, X. Kong, J. Sun, H. Yu, S. M. Smith, D. Wu and J. Li (2016). “Critical roles of soluble starch synthase SSIIIa and granule-bound starch synthase Waxy in synthesizing resistant starch in rice.” Proc Natl Acad Sci U S A 113(45): 12844–12849.

Zhou, W., J. Yang, Y. Hong, G. Liu, J. Zheng, Z. Gu and P. Zhang (2015). “Impact of amylose content on starch physicochemical properties in transgenic sweet potato.” Carbohydr Polym 122: 417–427.

Zhou, W., S. Zhao, S. He, Q. Ma, X. Lu, X. Hao, H. Wang, J. Yang and P. Zhang (2020). “Production of very-high-amylose cassava by post-transcriptional silencing of branching enzyme genes.” J Integr Plant Biol 62(6): 832–846.

Zhu, F. (2015). “Composition, structure, physicochemical properties, and modifications of cassava starch.” Carbohydr. Polym 122: 456–480.

